# Evidence of prezygotic isolation, but not assortative mating, between locally adapted populations of *Fundulus heteroclitus* across a salinity gradient

**DOI:** 10.1101/2022.09.03.506458

**Authors:** Reid S. Brennan, Andrew Whitehead

**Affiliations:** Marine Evolutionary Ecology, GEOMAR Helmholtz Centre for Ocean Research Kiel, Kiel, Germany; Faculty of Mathematics and Natural Sciences, Christian-Albrechts University of Kiel, Kiel, Germany; Department of Environmental Toxicology, University of California-Davis, Davis, CA, USA

**Author notes:** **Ethics statement** Experimental methods and animal care procedures were approved by University of California Davis IACUC protocol No.17221.

**Keywords:** Reproductive isolation, salinity, adaptation, fishes

## Abstract

Selection along environmental gradients can drive reproductive isolation and speciation. Among fishes, salinity is a major factor limiting species distributions and, despite its importance in generating species diversity, transitions between marine and freshwater are rare. Here, we tested for mechanisms of reproductive isolation between locally adapted freshwater and brackish water-native populations of killifish, *Fundulus heteroclitus*, from either side of a hybrid zone along a salinity gradient. There was evidence for pre-zygotic endogenous reproductive isolation with reduced fertilization success between crosses of freshwater-native males and brackish water-native females. Exogenous pre-zygotic isolation was also present where females had highest fertilization in their native salinity. We used a replicated mass spawning design to test for mate choice in both brackish and fresh water. After genotyping 187 parents and 2,523 offspring at 2,347 SNPs across the genome, 85% of offspring were successfully assign to their parents. However, no reinforcing mate choice was observed. These results therefore demonstrate emerging, yet limited, reproductive isolation and incipient speciation across a marine to freshwater salinity gradient and suggest that both endogenous and exogenous mechanisms, but not assortative mating, contribute to divergence.

## Introduction

Natural selection in divergent environments can result in local adaptation and ultimately drive speciation if barriers to gene flow emerge. During this process, ecological speciation (Schluter, 2000; Nosil, 2012), selection results in the fixation of alleles that are beneficial under one environment but not the other (Schluter, 2009). This in turn can drive reproductive isolation when it limits migration or when hybrids have reduced fitness in either environment. Thus, an essential aspect in understanding ecological speciation is how selection and genetic divergence facilitates reproductive isolation and speciation between populations in distinct environments.

Hybrid zones, where distinct populations or species meet, reproduce, and produce offspring (Barton & Hewitt, 1985), are particularly useful for studying the mechanisms that enable ecological speciation. When a locally adapted population transitions to another along an environmental gradient, gene flow between can be limited. Dispersal of individuals along the gradient may be constrained by selection against hybrids due to endogenous (intrinsic) genetic incompatibilities (Barton & Hewitt, 1989), thereby forming a “tension zone”. Conversely, hybrid zones can be maintained by environmental or exogenous (extrinsic) mechanisms, where locally adapted types have reduced fitness in the alternate environment (Hewitt, 1988).

To distinguish the mechanisms that limit gene flow between adjacent populations and potentially promote ecological speciation, it is necessary to understand the relative influence of exogenous and endogenous mechanisms. These can each take the form of prezygotic or postzygotic barriers. Prezygotic barriers include mechanisms such as mate choice, temporal isolation, and gametic competition, while postzygotic mechanisms may include hybrid inviability and reduced hybrid fitness (Coyne, 1992; Jiggins & Mallet, 2000). More specifically, endogenous prezygotic mechanisms are those that are independent of the environment yet occur before a zygote can be formed, such as mate choice. Prezygotic isolation can also be exogenous, for example if fertilization depends on specific environmental conditions. Similarly, postzygotic isolation may be endogenous such as when hybrids are always less fit than parents, but exogenous when hybrid fitness depends on the environment.

Environmental salinity provides a strong barrier to migration for aquatic species, where communities in adjacent fresh and salty habitats are almost entirely unique. That is, it is extremely rare for individuals from the same species to occupy both salty and fresh habitats, and this is true across the tree of life, including in fishes (Lee & Bell, 1999; Schultz & McCormick, 2012). Therefore, typically for a taxon to span marine and freshwater environments adaptive differentiation and speciation is required. A few micro-evolutionary model systems have provided insight into the mechanisms that enable differentiation across the marine-freshwater boundary. For example, in threespine stickleback (*Gasterosteus aculeatus*), selection in freshwater populations has driven divergence in armor plating (Bell, 2001), behavior (Di-Poi *et al*., 2014), body shape (Walker & Bell, 2000), salinity tolerance (Heuts, 1946), and diet (Ishikawa *et al*., 2019), among other traits (McKinnon & Rundle, 2002), compared to their marine ancestors. In most cases freshwater divergence is accompanied by reproductive isolation (McKinnon *et al*., 2004). Similarly, populations of the killifish *Lucania parva* are locally adapted to different osmotic environments (Kozak *et al*., 2014) and reproductive isolation has emerged across environmental salinity gradients (Kozak *et al*., 2012). Similar mechanisms likely maintain reproductive isolation between sister species *L. parva* (brackish/marine) and *L. goodei* (freshwater) (Kozak *et al*., 2015). Additional species such as European flounder (Momigliano *et al*., 2017), pike (Sunde *et al*., 2018), sockeye salmon (Wood & Foote, 1996), and others (Schluter, 1996), show patterns of divergence into low salinity and suggest that when a population is able to establish in freshwater, reproductive isolation and divergence can rapidly emerge.

Since most large fish clades are exclusively marine or exclusively freshwater (Betancur-R *et al*., 2015), micro-evolutionary model systems for studying the early/intermediate stages of differentiation along salinity gradients are rare. In addition to a few other models (e.g., sticklebacks, *Lucania* killifish), the Atlantic killifish *Fundulus heteroclitus* provides a model system to study this process. *F. heteroclitus* is a euryhaline species distributed along the eastern coast of North America, from Florida to Nova Scotia, and across the entire salinity continuum from marine to freshwater (Hildebrand & Schroeder, 1928). At least two distinct hybrid zones delineate adaptive differentiation across two different environmental gradients across the range. The first hybrid zone is found in coastal New Jersey, and distinguishes northern and southern coastal populations (sub-species) (Ropson *et al*., 1990; McKenzie *et al*., 2016), where sub-species differ in genetics, morphology (Able & Felley, 1986), and thermal physiology (DiMichele & Powers, 1982; Fangue *et al*., 2006). Individuals from either side of the hybrid zone (New Hampshire and North Carolina) showed evidence of endogenous isolation in the form of reduced hatching success as well as assortative mating (McKenzie *et al*., 2017).

In addition to the coastal zone, another hybrid zone along a salinity gradient is found within the Chesapeake Bay (Whitehead *et al*., 2011). While ancestrally marine, *F. heteroclitus* has broad osmoregulatory flexibility (plasticity) and can tolerate a wide range of salinities from dilute freshwater to four-times the salinity of seawater (Griffith, 1974; Whitehead, 2010). Individuals native to freshwater (FW-native) are genetically distinct from their downstream brackish counterparts (BW-native) with a genetic cline centered on the tidal freshwater boundary (∼1/32^nd^ the salinity of seawater; ∼1 parts per thousand, ppt) (Duvernell *et al*., 2008; Whitehead *et al*., 2011). This 1 ppt salinity is also physiologically important. Acclimation to salinities across 1 ppt (above or below) requires extensive remodeling of transport epithelia (e.g., gills) (Whitehead *et al*., 2012) which consumes ∼10% of their energy budget (Kidder *et al*., 2006a; b). Populations native to each salinity diverge in their salinity-specific performance, for example showing differences in transcriptional and osmoregulatory plasticity (Whitehead *et al*., 2011; Brennan *et al*., 2015), divergence in salinity specific swimming performance (Brennan *et al*., 2016), and adaptive genetic divergence due to selection (Brennan *et al*., 2018). Thus, fresh- and brackish-native populations are locally adapted to their respective osmotic environments and remain genetically distinct despite the presence of a narrow hybrid zone populated by highly admixed individuals (Brennan *et al*., 2018).

While there is much evidence that the FW-native and BW-native populations are adaptively diverging across this salinity gradient, the mechanisms that are preventing the homogenization of the populations and maintaining the hybrid zone are not known. Previous work has suggested that hybrids may have reduced low-salinity tolerance relative to the parental populations (Brennan *et al*., 2018), but no studies have yet attempted to identify the mechanisms contributing to reproductive isolation between brackish and freshwater populations.

Here, we test for the presence of reproductive isolation that may serve to constrain gene flow between FW-native and BW-native populations of *F. heteroclitus* across a salinity gradient. We ask whether endogenous and exogenous prezygotic and postzygotic isolation exist, which may limit gene flow between the populations. Given the coincidence of the hybrid zone with a physiologically stressful environmental salinity we predicted that environmental salinity would affect reproductive success between the populations. To test this, we quantified reproductive incompatibilities between FW-native and BW-native parents, and mate choice, in both fresh water and brackish water. First, manual crosses were used to determine if populations could successfully interbreed and produce viable offspring. Second, mate choice was assessed using a mass spawning design where individuals were free to interbreed over the course of a month. We genotyped 2,523 offspring and 187 parents at 2,347 single nucleotide polymorphisms (SNPs) across the genome using a RAD-capture approach (Rapture; (Ali *et al*., 2016), and assigned parentage to > 2,000 of these offspring to determine the mate choice of all parents. These two experiments were conducted in both fresh and brackish water.

## Methods

Adult *F. heteroclitus* were collected in September, 2014 and March 2016. FW-native fish were sampled from the Potomac River at Piscataway Park, near Accokeek, Maryland, USA (38°41′42.18′′N, 77°3′10.38′′W); salinity has not been higher than 0.12 ppt since monitoring began in 1986 (chesapeakebay.net). BW-native fish were from Point Lookout State Park, Scotland, Maryland, USA (38°3′10.90′′N, 76°19′34.38′′W), where the average salinity is approximately 11.8 ppt. Fish were shipped overnight to the University of California Davis and held in 30-gallon tanks at the salinity close to their collection salinity in 850 gallon recirculating systems equipped with UV, mechanical, and biological filters. Fish were kept at a density less than one per gallon, fed Aquaxcel 4512 daily (Cargill, Minnesota, USA). The light cycle was 10 h light and 14 h dark with 21°C water temperature. Dechlorinated tap water was used for freshwater (0.2 ppt) and reverse osmosis water mixed with Instant Ocean Sea Salt (Instant Ocean, Spectrum Brands, VA, USA) was used to achieve brackish salinity (15 ppt). Fish were randomly transferred to their experimental salinity and allowed to acclimate for at least one month. To distinguish the populations when combined in a single tank, all fish were marked with a fluorescent elastomer tag (Northwest Marine Technology, WA, USA).

To test for postzygotic isolation, we used five replicate manual crosses for each of four cross types: two within-population (BW-native x BW-native, FW-native x FW-native) and two between-population (BW-native♂ x FW-native♀, FW-native♂ x BW-native♀). These 20 crosses were repeated in two salinities: fresh water (0.2 ppt) and brackish water (15 ppt) (40 total crosses). For each replicate, three males and three females were combined in 30-gallon tanks with one 10.2 cm diameter spawning basket and allowed to freely spawn for 30 days. These baskets were covered with 9.5 mm mesh through which spawned embryos could easily fall but adults could not gain access. This is necessary as adults will consume eggs and embryos. Preliminary trials showed that adults recognize these baskets as the primary spawning substrate in the tanks. Baskets were checked for eggs 3 times per week. All eggs were transferred to 100 × 15 mm plastic petri dishes and checked for fertilization under a stereomicroscope. Fertilization was determined by the appearance of the perivitelline space, which develops minutes after fertilization, or the neural keel, which is visible ∼40 hours post fertilization (Armstrong & Child, 1965). Fertilized eggs were incubated in 3 ppm methylene blue (Kordon) to prevent fungal growth and water was changed daily. The number of total eggs, fertilized eggs, and hatched fry were recorded. Typical time to hatching is 14 days (Armstrong & Child, 1965), and embryos that did not hatch after 21 days were considered inviable. The brackish water trial was conducted from May 14 to June 12, 2016 and the freshwater trial was conducted from August 14 to Sept 7, 2016.

Fertilization and hatching success were analyzed using logistic regressions in R version 4.1.2 (R Core Team, 2022). The response variable was either hatching success counts or fertilization success counts, and main effects included salinity environment (hereafter referred to as “salinity”) and cross type (hereafter referred to as “cross”), and their interaction. We tested for the significance of main effects with wald tests using aod version 1.1 (Lesnoff & Lancelot, 2012) and post hoc differences were calculated with lsmeans version 2.30 (Lenth, 2016).

Mate choice was investigated using a “choice test” in a free spawning design. In each of four replicate 30-gallon tanks at both freshwater (0.2 ppt) and five replicate tanks at brackish water (15 ppt) (9 total tanks), five males and five females from both the FW-native and BW-native populations (20 individuals per tank) were combined. Two spawning baskets were placed on either end of each tank to collect eggs. Three replicates of within population controls were used per population to ensure that both populations were reproductively active. These controls contained three males and three females from one population and one spawning basket. Eggs were collected every two days, checked for fertilization, and incubated as described in the no-choice test. For all trials, dead fish were replaced with an individual of the same sex and population (6 total deaths). Upon hatching, fry were preserved for genotyping in order to assign parentage and identify the cross-type from which they were derived. All samples were stored at -80°C until processing. Unhatched eggs were again considered non-viable after 21 days of development.

### Genotyping

Genomic DNA was extracted from whole fry or fin clips with a custom method as in Brennan et al. (2018). Between-population crosses resulted in unequal numbers of hatched fry and replicates with >350 hatched individuals, fry were subsampled down to 350 samples. This included BW replicate 1, 3, and 4, which were reduced to 41%, 45%, and 75% of the total individuals, respectively.

A restriction-associated DNA sequencing (RAD-seq) capture approach (Rapture) was used for genotyping as described by Ali et al (2016). The strength of this approach is that it allows high-throughput and low-cost genotyping of thousands of samples at hundreds to thousands of loci. 250 ng DNA from each sample was digested with the restriction enzyme *Sbf*I. In sets of 96, individuals were barcoded by adding 2 *μ*l indexed SbfI biotinylated RAD adapter (50 nM), each with a unique 8-bp barcodes. DNA was sheared to 200-500 bp using a Bioruptor NGS sonicator, RAD-tagged DNA fragments were isolated, and the final DNA was prepared for Illumina sequencing using a NEBNext Ultra DNA Library Prep Kit for Illumina. A unique barcode was applied to each set of 96 individuals, enabling multiplexing of thousands of individually-indexed individuals.

We designed 866 capture baits based on previously collected *Sbf*I RAD data (Brennan *et al*., 2018), specifically targeting loci that contained known polymorphisms between the two populations at intermediate frequencies. These baits were also chosen based on their unique location in the reference genome (on different scaffolds) (Reid *et al*., 2017), unique sequence composition, and optimal GC content (see Data availability for more details). Capture baits (MYbaits) were ordered from MYcroarray (currently Arbor Biosciences, MI, USA). This baitset captured 103,920 bases, plus flanking regions. Equal amounts of all libraries were pooled to a final concentration of 30 ng/*μ*l. This pool was split evenly between 3 independent capture reactions. Sequence capture proceeded according to the manufacturer’s protocol except for the primers used in the final amplification step. Here we substituted universal primers of the sequence AATGATACGGCGACCACCGAGATCTACACTCTTTCCCTACACGAC*G and CAAGCAGAAGACGGCATACG*A, where * is a phosphorothioated DNA base.

Libraries were sequenced using 150-bp paired-end reads on an Illumina Hiseq 4000 at the UC Davis Genome Center. Reads were demultiplexed first by Illumina barcode and then by RAD barcode. For RAD barcodes, perfect barcode matches with a partial restriction site were required. Reads were aligned to the reference genome version GCF_000826765.1 (Reid *et al*., 2017) using BWA-MEM version 0.7.12 (Li, 2013) and PCR duplicates were removed using SAMBLASTER (Faust & Hall, 2014). We removed 117 offspring with mean coverage < 5x. Next, variants were filtered to include sites with minor allele frequency greater than 0.05, only biallelic sites, depth between 10 and 110x, minimum quality of 20, and data in at least 90% of individuals. Finally, this set was filtered to include only high confidence SNPs in the parents (to ensure accurate parentage assignment). Across all parental individuals, minor allele frequency was required to be at least 0.01 with at least 85% of individuals with non-missing data. From this, 2,347 SNPs were obtained.

To visualize population structure between parental individuals, a principal component analysis was run on the parent samples only using PLINK2 (Chang *et al*., 2015) with variants pruned for linkage disequilibrium (LD; 1,606 variants remained). Weir and Cockerham’s Fst (Weir & Cockerham, 1984) was estimated using VCFtools version 0.1.15 (Danecek *et al*., 2011).

Parentage assignment was conducted using Colony v. 2.0.6.6 (Jones & Wang, 2010). Within each replicate, SNPs were LD pruned and parentage was assigned to the known list of candidate parents using the pairwise likelihood approach. Offspring were filtered to only those that had two parents assigned with 95% confidence. With these data, we were unable to determine the actual number of independent mating events because multiple matings between the same two individuals would be undetectable. Therefore, we considered the mate choice as the proportion of within-population versus between-population mate pairings for each individual, as inferred from parentage assignment of offspring. For example, if one male parent from the BW population produced offspring with one BW female and three FW females, the proportion of like-type matings would be 25%. To test for mate choice, data were analyzed with a logistic regression. The count of like versus unlike mate pairings was considered the response variable, whereas main effects included treatment salinity (environment), source population, and sex (and their interactions). Individual length was treated as a covariate, and tank was treated as a random effect. Neither length or tank improved model fit and were dropped from the final models. Models were run using the *glm* function in lme4 (Bates *et al*., 2014). *Wald tests* were used to test for a statistically significant influence of main effects, with post hoc tests using *lsmeans (Lenth, 2016)*, and the *effects* package was used to visualize model predictions (Fox, 2003; Fox & Weisberg, 2018a; b).

## Results

### Population structure

Principal components analysis (PCA) of Rapture SNP variation among 186 adult fish showed that the parental populations native to different osmotic environments were genetically distinct (Fig. 1). Weir and Cockerham’s weighted FST between populations was 0.077 which is consistent with other estimates (RAD-seq SNPs: 0.082 (Brennan *et al*., 2018); Microsatellites: 0.081 (Whitehead *et al*., 2011)). ADMIXTURE estimates of individual ancestry similarly support the distinctiveness of the parental populations and also demonstrate that all offspring follow the expected ancestry given their parental assignment (Fig. S1; details of parentage analysis below) (Alexander *et al*., 2009).

**Figure 1:**
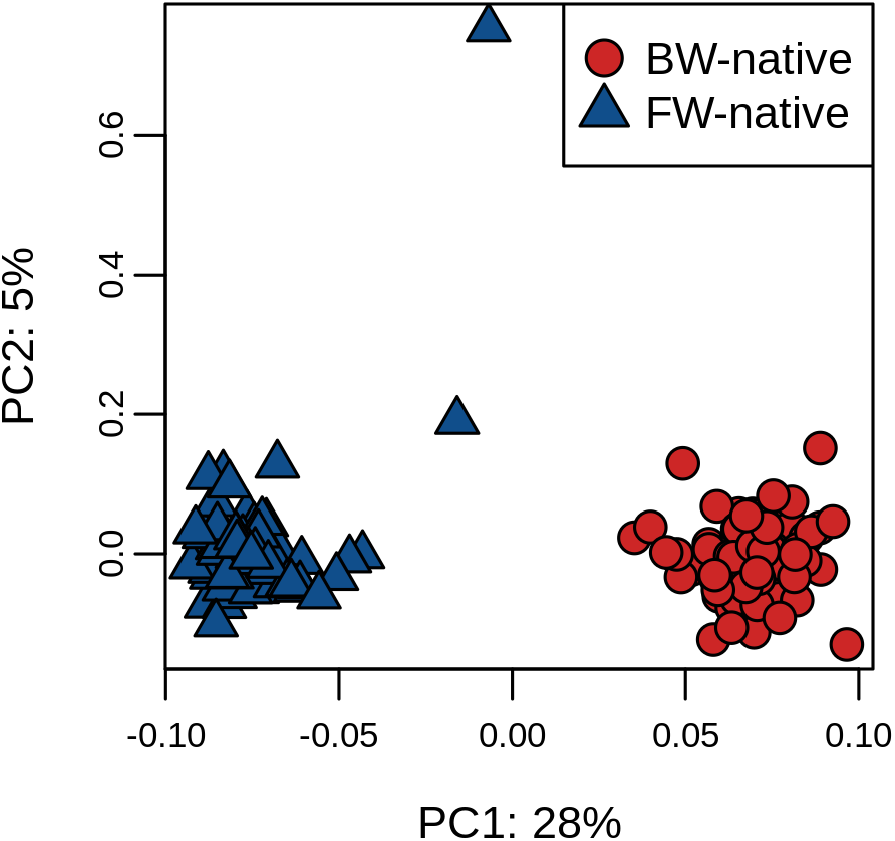
Principal components analysis of 1,606 linkage disequilibrium pruned SNPs for all BW-native (red circles) and FW-native (blue triangles) parents used in the mate choice experiment. Axis labels show the percent of the total variance explained by principal components one and two.

### Reproductive incompatibilities

Fertilization and hatching success were high across all cross-type and salinity environment treatment groups during the tests for reproductive incompatibilities (mean ± standard error; fertilization = 0.90 ± 0.01; hatching = 0.84 ± 0.02). Variation in fertilization success between populations and between environments provided evidence of endogenous and exogenous reproductive isolation (Fig. 2A). For endogenous isolation, there was an effect of cross (Wald test, *χ*^2^(3)=32.2, P = 4.8e-07) where crosses between FW-native♂ and BW-native♀ had lower fertilization success than all other crosses (P < 0.05), though the effect size of this was small (maximum reduction: 8.7%; reduction of fertilization success ± std. error: 0.082 ± 0.0126). While there was no significant effect of salinity (*χ*^2^(1)=2.9, P = 0.086), the marginally significant interaction between salinity and cross (*χ*^2^(3)=7.8, P = 0.05) suggests the possible influence of extrinsic isolation due to salinity environment. This interaction effect was due to the lower fertilization success of BW-native♀ in fresh water (0.831 ± 0.019; 0.912 ± 0.009) relative to brackish (0.875 ± 0.008; 0.931 ± 0.006) as compared to higher success of FW-native♀ in fresh water (0.949 ± 0.01022; 0.910 ± 0.02302) relative to brackish (0.920 ± 0.0129; 0.900 ± 0.0245; Fig. 2A). That is, females tended to have higher fertilization success in their matched salinity environment (e.g., BW-native♀ in BW environment, and FW-native♀ in FW environment) compared to their un-matched salinity environment (e.g., BW-native♀ in FW environment, and FW-native♀ in BW environment).

**Figure 2:**
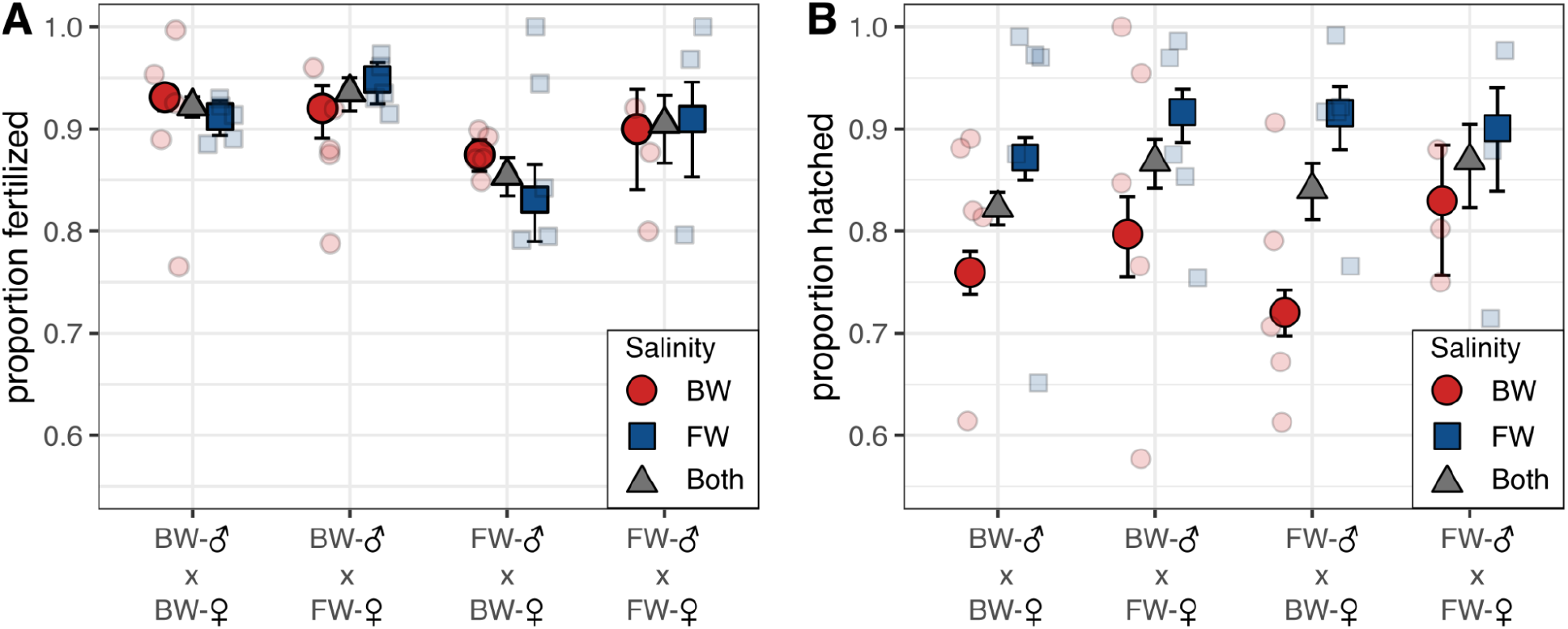
Results from the manual crosses for (A) fertilization and (B) hatching success. Small transparent points show individual replicates while larger bold colored points are the model estimates including standard error. Results are separated by salinity where red circles are the brackish treatment and blue squares are the freshwater treatment. The gray triangles are the average of both salinities within the cross type. (A) The FW-native♂ x BW-native♀ cross had lower fertilization success than other crosses and there was an interaction effect where BW-native♀ in brackish water had higher fertilization than in fresh water. The opposite pattern was detected for FW-native♀ with had higher fertilization success in fresh water than in brackish water. (B) Hatching success showed a significant salinity effect and was higher in fresh water. There was also a significant effect of cross where BW-native♂ x BW-native♀ had lower hatching than BW-native♂ x FW-native♀. There was a significant interaction between cross and salinity due to a significant salinity effect in all cross types except FW-native♂ x FW-native♀ crosses.

For hatching success we detected a significant effect of salinity (*χ*^2^(1)=46.9, P = 7.5e-12) with consistently higher hatching in fresh versus brackish water across crosses (difference ± std. error: 0.123 ± 0.0146) (Fig. 2B). But in contrast to fertilization, there was no evidence of reduced hatching success for inter-population crosses relative to within-population crosses (P > 0.05). There was a significant effect of cross-type (*χ*^2^(3)=17.1, P = 0.0007) where BW-native♂ x BW-native♀ had lower hatching success than BW-native♂ x FW-native♀ (difference ± std. error: 0.045 ± 0.015), though this was a relatively small effect size (5% reduction). Finally, there was a significant interaction between cross and salinity (*χ*^2^(3)=8.8, P = 0.032) that was driven by differences between brackish and freshwater treatments that was apparent for all crosses (P < 0.0001) except for FW-native♂ x FW-native♀ (P = 0.67).

### Mate choice

For the control parental tanks and the experimental tanks, we observed high levels of both fertilization (mean ± standard error: FW-Native: 0.83 ± 0.04; BW-Native: 0.85 ± 0.02; Experimental tank: 0.92 ± 0.01) and hatching success (FW-Native: 0.76 ± 0.06; BW-Native: 0.74 ± 0.06; Experimental tank: 0.74 ± 0.03). If there were differences in fertilization success or hatching success between the populations we might have observed apparent differences in mate choice that were actually due to physiological limitations. The successful reproduction of each parental population provides confidence that the observed differences in mate pairings were due to choice rather than physiological limitations. For mate choice, complete data were collected for nine experimental tanks: five replicates in the brackish salinity environment and four replicates in the freshwater salinity environment. One FW environment replicate was lost because BW-native males were accidentally not added to this treatment. An average of 433 ± 84 eggs were produced per experimental replicate (14.4 eggs/day) with a maximum of 1,431 and a minimum of 142.

Overall, 2,523 offspring with greater than 5x sequence coverage (Fig. S2) and 187 parents, were genotyped at 2,347 SNPs across the genome. Of the 2,523 offspring, 85% (2,138) were successfully assigned to two parents each with 95% confidence. Overall, despite removing individuals with coverage < 5x, low parentage assignment was in excess for individuals with low sequencing depth, but it was not restricted to only low depth individuals (Fig. S3). All samples and parents were also run in a single Colony analysis and the number of impossible parents (i.e., those not in the same experimental tank) was determined. Only 1% of assignments (25) were impossible (assigned to parents from different tanks), providing confidence that the parentage assignment was robust.

Of the 187 parents, 137 had at least one offspring assigned, representing 63 males and 74 females. The number of offspring assigned to each individual was variable and ranged from 1 to 201 (mean ± standard error: 31.2 ± 3.5) and was similar between males (33.9 ± 5.9) and females (28.9 ± 4.0; two-sided t-test: P = 0.48; Fig. S4). Overall, family size ranged from 1 to 89 with a mean of 11 ± 1.15. Finally, the number of mates per individual with assigned offspring ranged from 1 to 9 (2.77 ± 0.16) with no difference between males (3.02 ± 0.27) and females (2.57 ± 0.17; two-sided t-test: P = 0.17) and no relationship with the length of the individuals (ANOVA: P = 0.99; Figs. S6).

There was limited evidence of mate choice. A random effect of experimental replicate did not improve model fit (Likelihood ratio test: P = 1) and was not included in the final model. The mean size of adults was 7.04 ± 0.05 cm and FW-native individuals (7.26 ± 0.072 cm) were significantly larger than BW-native individuals (6.85 ± 0.06 cm; P = 0.0003; Fig. S7). Despite this, there was no effect of length on mate choice (P = 0.99; Fig. S8). Overall, the results of the logistic regression did not support the influence of mate choice in these data (Fig. 3) as there was no main effect of salinity (Wald test, *χ*^2^(1)=0.004, P= 0.98), cross (*χ*^2^(3) = 5.5, P=0.14), or their interaction (*χ*^2^(3)=5.8, P=0.12). Consistent with this, all model estimates overlapped and most, with the exception of BW-native females in FW (estimate and 95% CI: 0.68 [0.51, 0.81]), were not different from 0.5 (Fig. 3), indicating that mating was largely random throughout the experiment.

**Figure 3:**
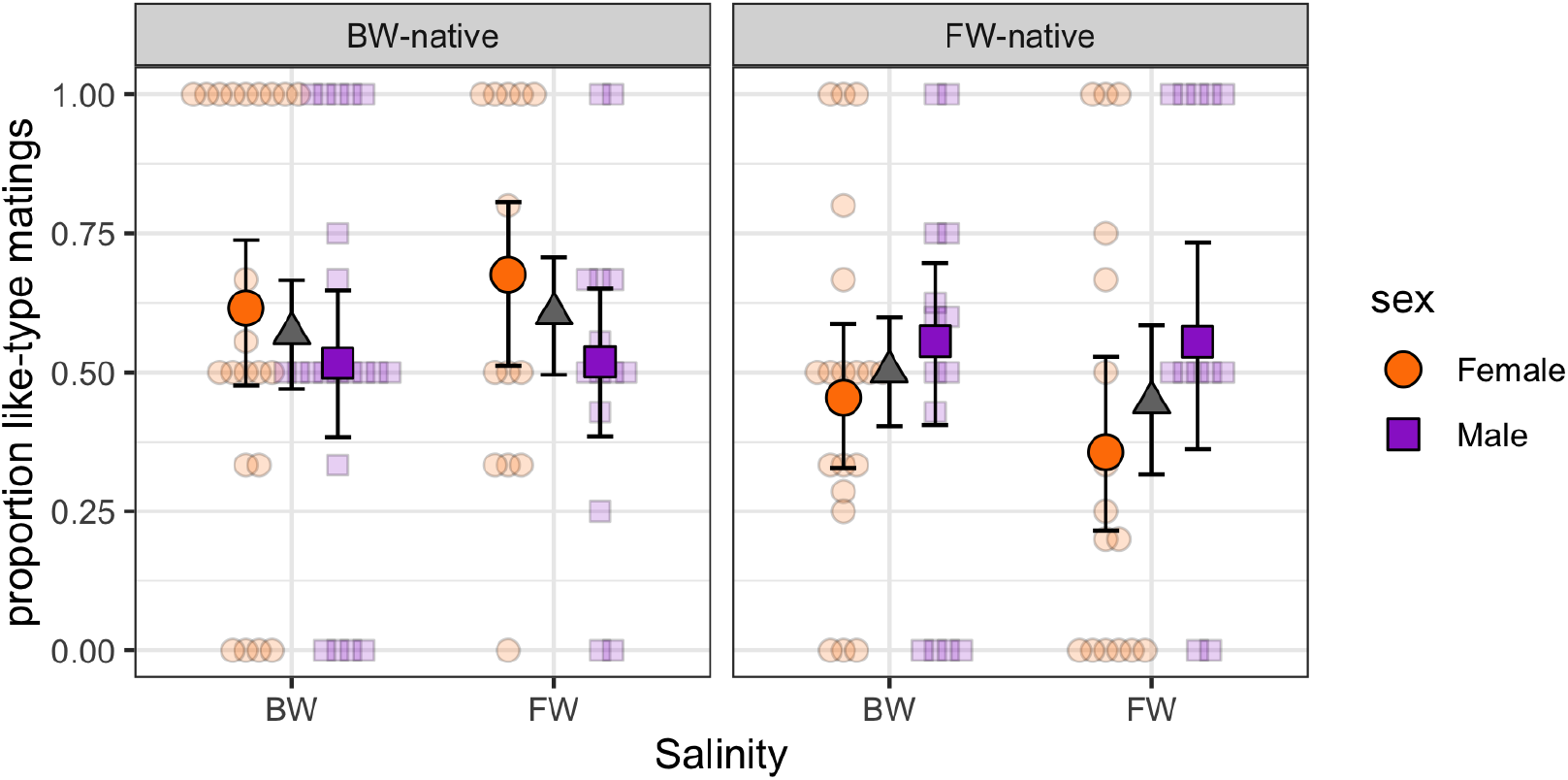
Results from the mate choice test for assortative mating. The y-axis is the proportion of pairings with mates from the same population (like-type matings). The left panel shows BW-native individuals and the right panel FW-native individuals. Within each panel points are separated by the environmental salinity, and the color and shape of each point represents the sex (orange circles for female and purple squares for male). Small transparent points are the results from each individual replicate while larger bold-colored points are the model estimates for the treatment with standard error. The gray triangles are the average across both sexes within a treatment. There were no significant main effects or interactions.

## Discussion

*Fundulus heteroclitus* are distributed across steep salinity gradients where there is strong genetic structure, local adaptation to alternate osmotic environments, and reduced performance of hybrids. An apparent barrier to gene flow is localized to a region of the salinity gradient that is physiologically important (∼1 ppt), and co-localizes with a zone of admixture (hybrid zone). As such, we hypothesized that exogenous isolating mechanisms were restricting gene flow between populations. Indeed, we detected evidence for prezygotic exogenous isolation, where females tended to have higher fertilization success in their matched salinity environment, compared to when fertilizations were attempted in their unmatched salinity environment (Fig. 2A).

Simultaneously, we detected prezygotic endogenous isolation, where FW-native♂ x BW-native♀ hybrids had reduced fertilization success regardless of salinity. No evidence of postzygotic isolation was found (Fig. 2B). While the effect sizes were relatively small, these mechanisms could contribute to reduced gene flow near the tidal freshwater boundary that distinguishes populations inhabiting fresh versus salty water.

Salinity tolerance of gametes has a major impact on the ability of a population to occupy novel osmotic environments (Green *et al*., 2020) and mismatch between gametes and external salinity can limit gene flow even if adults can survive. If there is a mismatch between the necessary activation signal or conditions for gametes to stay active, successful reproduction can be greatly reduced, known as immigrant reproductive dysfunction (IRD) (Svensson *et al*., 2017). IRD can limit gene flow between populations and facilitate ecological speciation. For instance, sand goby, *Pomatoschistus minutus*, and Atlantic herring, *Clupea harengus*, both show divergence in sperm motility between populations locally adapted to different salinities (Berg *et al*., 2019; Leder *et al*., 2021). Further, salinity specific sperm motility is common between closely related species that occupy different osmotic environments (Elofsson *et al*., 2003; Lindström *et al*., 2021). Our results support these previous studies and show that low salinity drives a small reduction in fertilization success for BW-native♀ (Fig. 2). Thus, IRD may be promoting divergence in this system. However, it should be noted that this is a conservative estimation as there are numerous environmental differences between the two habitats (e.g., pH, predators, nutrients, pathogens, oxygen, food, symbionts, tidal rhymicity, etc.) that could influence performance and fitness between the populations and this limit gene flow (Austin & Austin, 2007; Lozupone & Knight, 2007; Osborne *et al*., 2015; Ou *et al*., 2015).

To test for the influence of assortative mating, we leveraged high-throughput genotyping with a replicated mass spawning experiment across multiple environments. We were able to accurately and rapidly genotype and assign parentage to thousands of offspring. This approach recapitulated known genetic relationships between the parental populations and resulted in high parentage assignment across the experiment. However, we did not detect evidence of assortative mating at either salinity (Fig. 3). This is in contrast to other model systems, such as cichlids or swordtails, where assortative mating and sexual selection is the dominant factor limiting homogenization of populations and species (Knight & Turner, 2004; Salzburger *et al*., 2006; Schumer *et al*., 2017). Similarly, in the closely related killifish, *Lucania sp*., strong assortative mating was identified between populations and sister species adapted to different salinities (Fuller, 2008; Berdan & Fuller, 2012; Kozak *et al*., 2015), but in none of these cases did assortative mating differ between salinities. The lack of mate choice in *F. heteroclitus* suggests that the sexual selection and assortative mating observed in other systems seems to play a more minor role. However, mate choice is generally impacted by the condition of individuals, where unhealthy individuals will tend to find fewer mates (Neff & Cargnelli, 2004). Thus, under natural conditions it is possible that failure of a non-native fish to completely acclimate to a mis-matched osmotic environment could impair long-term health and performance, thereby leading to mate choice and selection against migrants (Hendry, 2004). Additional studies would be required to further explore this hypothesis.

Although we detected the influence of pre-zygotic mechanisms that may contribute to limiting gene flow across the adaptive transition zone at the tidal freshwater boundary, the effect sizes were not large. Here we consider caveats to our experimental design, and the potential influence of other mechanisms. We treated developing embryos with fungicide, which is common practice in fish husbandry. However, research in *Lucania* killifish found that hybrids between FW and BW types had reduced survival in fresh water, but that this effect was masked when embryos were treated with fungicide (Kozak *et al*., 2012). Future experiments with *F. heteroclitus* may consider trials both with and without fungicide treatment. It is also possible that migrants and hybrids are less fit than native individuals inhabiting their native salinity environment but assortative mating has not emerged. Assortative mating is less efficient at maintaining hybrid zones than reductions in hybrid fitness; a small reduction in hybrid fitness will have a much larger effect than even strong mate preference (Irwin, 2020). We have previously shown that admixed individuals have reduced physiological tolerance to fresh water relative to either parental population, suggesting impaired hybrid fitness. This could limit gene flow and serve to maintain genetic differentiation between the locally-adapted populations inhabiting either side of the tidal freshwater boundary. Additionally, behavioral effects could be present in the wild that may not manifest in captivity. For example, evidence for assortative mating in swordfish under field conditions disappeared when animals were studied in captivity (Schumer *et al*., 2017). Egg deposition locations differ between coastal populations *F. heteroclitus* (Able, 1984). If mating events occur at different or unsuitable locations in each salinity environment, reproduction of immigrants could be reduced, limiting gene flow. Indeed, potential killifish spawning substrates (aquatic plants) differ between fresh and brackish water habitats (Haramis & Carter, 1983).

Previous work across coastal temperature gradients in *F. heteroclitus* has shown reduced hatching rates and mate choice between northern and southern populations (McKenzie *et al*., 2017). Given this, we expected to observe similar patterns of reproductive isolation between FW-native and BW-native populations. The lack of agreement could be due to lower genetic differentiation between populations that span the Chesapeake salinity transition zone (Fst = 0.077) than between those that span the coastal latitudinal transition zone (Fst = 0.180) (Adams *et al*., 2006). Greater time since isolation can lead to increased genetic differentiation and accumulation of incompatibilities (Coyne & Orr, 1989). Furthermore, the nature of natural selection and adaptation along the coastal gradient (thermal physiology) and the Chesapeake gradient (osmoregulatory physiology) differs and could therefore have different impacts on reproductive and developmental traits (Lindtke & Buerkle, 2015). Finally, experimental design considerations could play a role. For example, McKenzie et al. used strip spawning to generate a single experimental replicate with 47 individuals, whereas we used natural crosses across nine replicates each with 20 individuals.

The results presented here show the emergence of reproductive isolation between populations of killifish locally adapted to local salinity and indicate the potential initial steps of incipient speciation. Additional work could help disentangle the drivers underlying this emerging isolation. For example, identifying the mechanisms leading to the higher fertilization success of females in their native salinity could provide insight into how osmotic environments can limit gene flow. Further, behavioral assays, including spawning substrate preference, could provide additional nuance into how the populations interact and may give more detailed information as to the mating preferences of each population. Finally, it is necessary to understand the impact of other abiotic and biotic factors that differ between marine and freshwater environments on population performance and ultimately gene flow. By understanding the mechanisms that lead to reproductive isolation and speciation across salinity gradients, we can gain insight into the potential mechanisms that have enabled diversification of fishes between marine and freshwater environments.

## Acknowledgements

This work was supported by funding from the National Science Foundation (DEB-1265282 to A.W., and DEB-1601076 to R.S.B. and A.W.).

## Data availability

Raw sequence reads are available at the National Center for Biotechnology Information (NCBI; Bioproject no. PRJNA872341). Other data including hatching success, fertilization, mate choice, and the capture probe fasta file as well as code to run all analyses are available on Zenodo (https://doi.org/10.5281/zenodo.7041408).

