## Supplemental Figures for "Evidence of prezygotic isolation, but not assortative mating, between locally adapted populations of *Fundulus heteroclitus* across a salinity gradient"

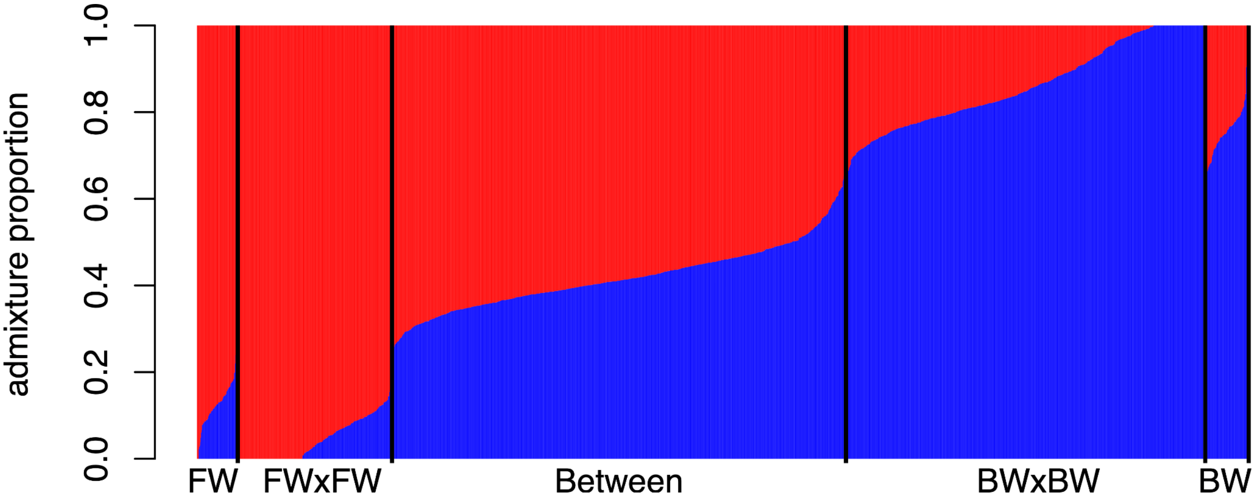


**Figure S1**: Admixture proportions calculated from Admixture. Each bar is an individual where the color indicates the assigned ancestry. Vertical black lines separate groups of individuals where "FW" are the FW-native parents, "BW" are the BW-native parents, "FWxFW" and "BWxBW" are the offspring assigned to parents of the same population, and "Between" indicates offspring with one parent from each population. All parentage assignments are from Colony as described in the main text.

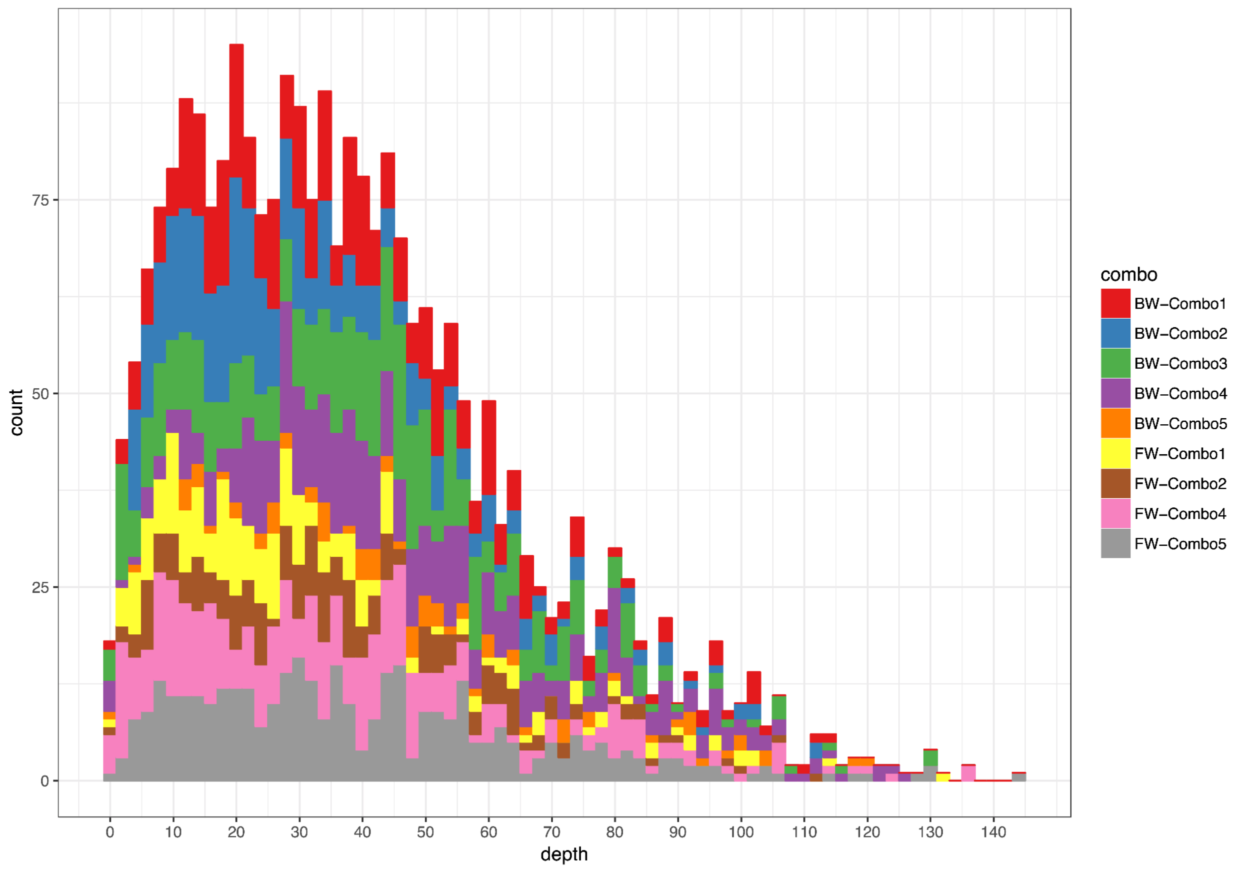


**Figure S2:** Histogram of average depth of sequencing for each individual. For each probe, the depth at the 20th base was determined and the mean calculated across all probes. The color indicates the experimental tank.

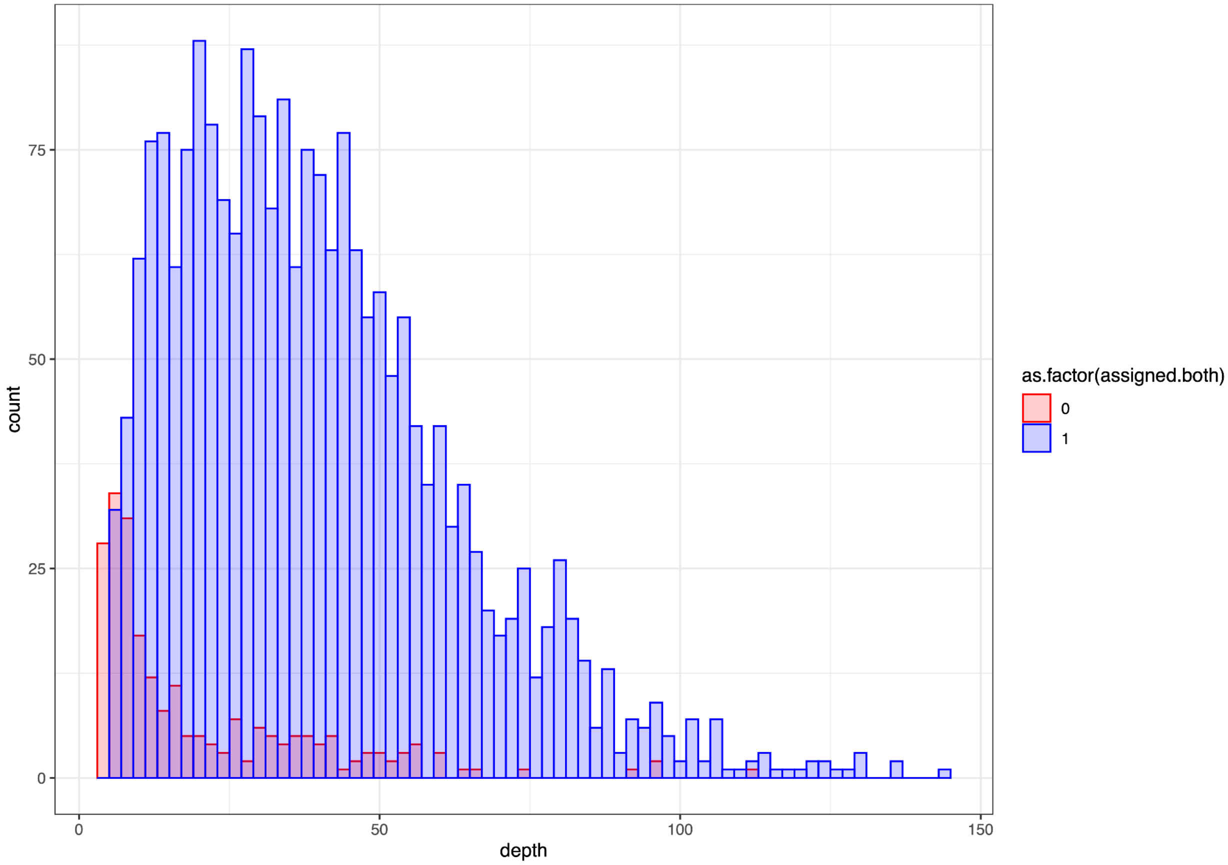


**Figure S3:** Assignment rate by sequencing depth as calculated in figure S2. Color indicates if an individual had both parents successfully assigned (blue) or not (red). Low coverage individuals were generally more likely to not be assigned parents successfully.

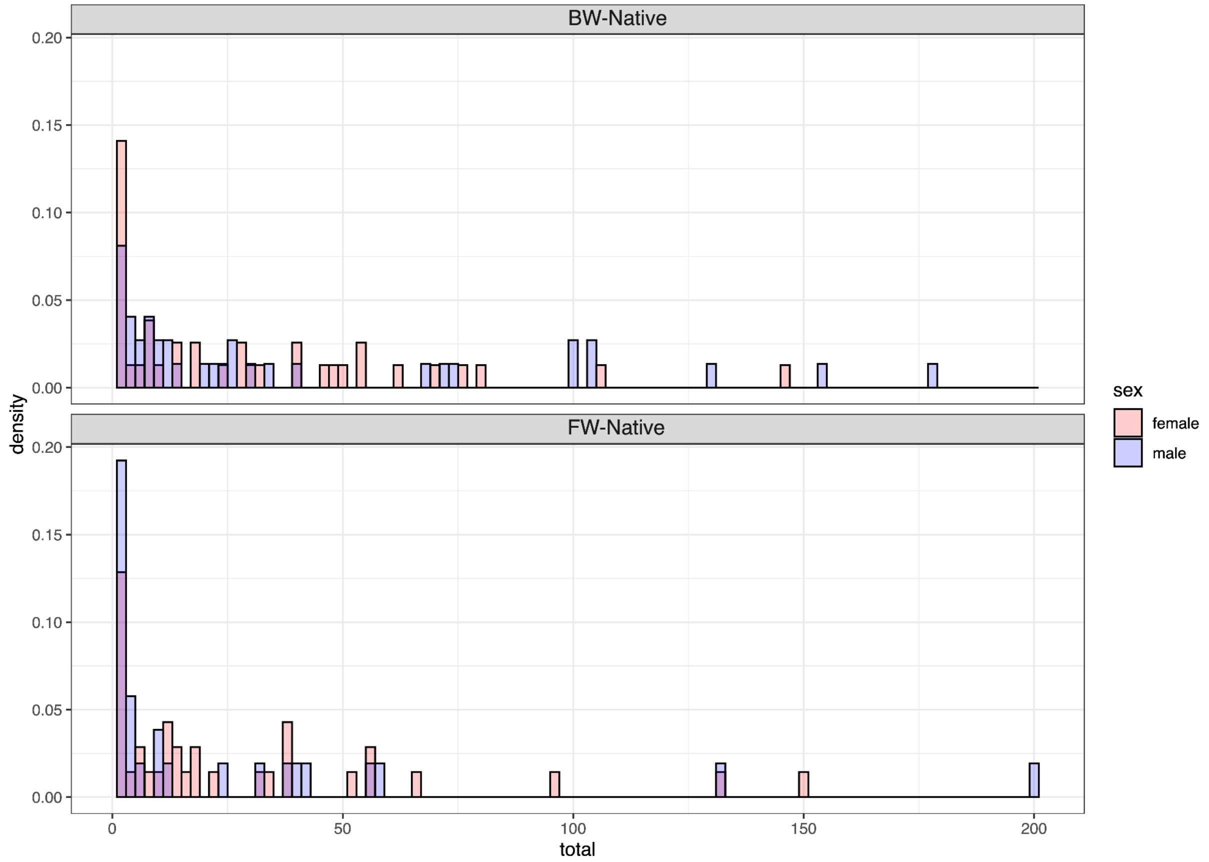


**Figure S4:** The number of offspring assigned to each parent. Color indicates sex of the individual.

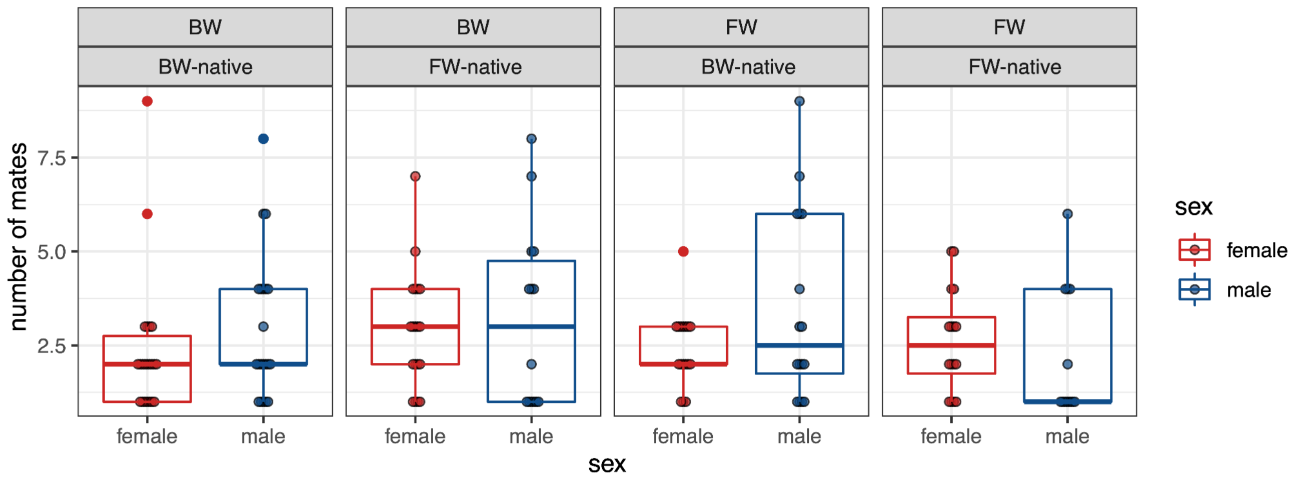


**Figure S5:** Number of mates for each individual with assigned offspring. The top label indicates salinity and the second label shows the population.

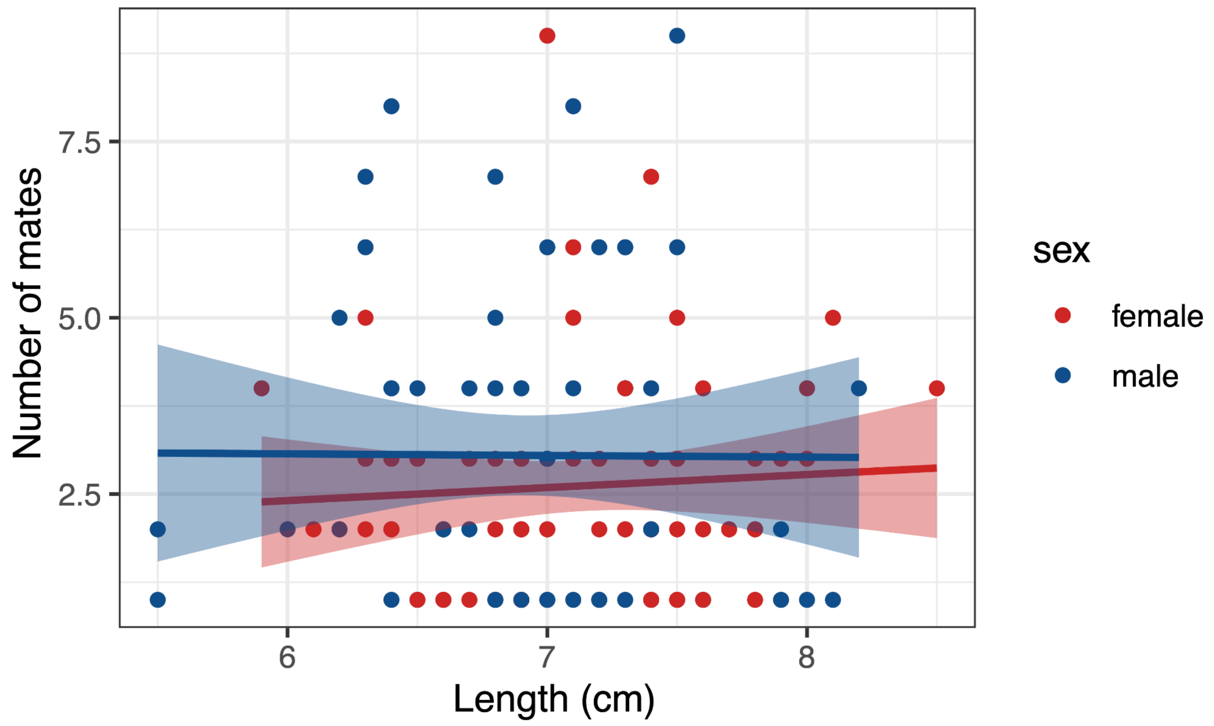


**Figure S6:** Relationship between adult size and the number of mates.

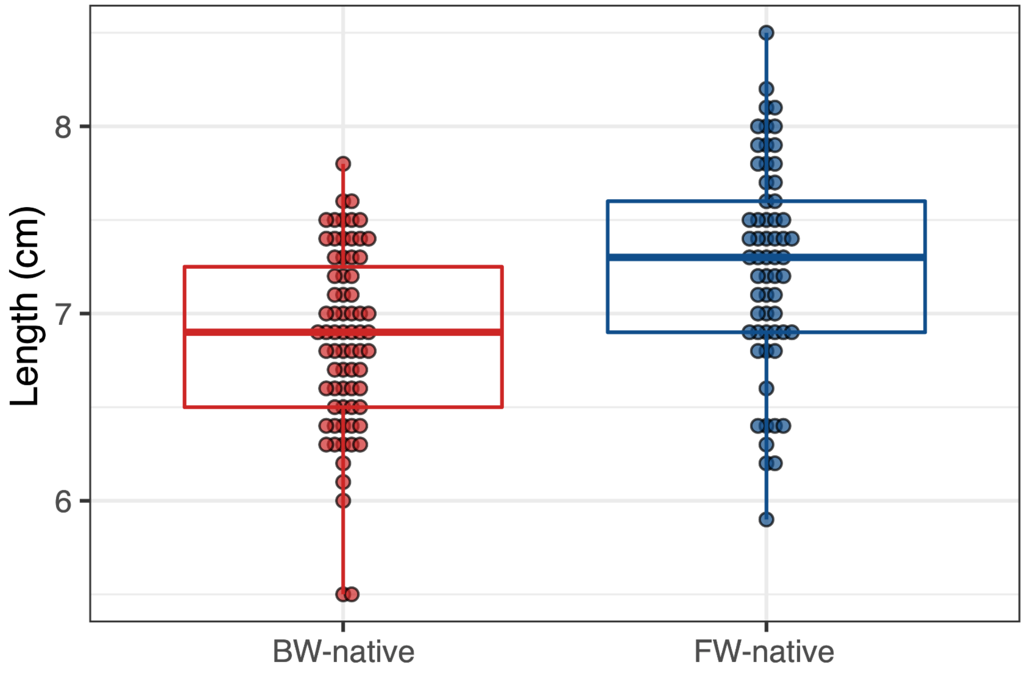


**Figure S7:** Length of parents used in the choice experiment. FW-Native individuals were significantly larger than BW-native.

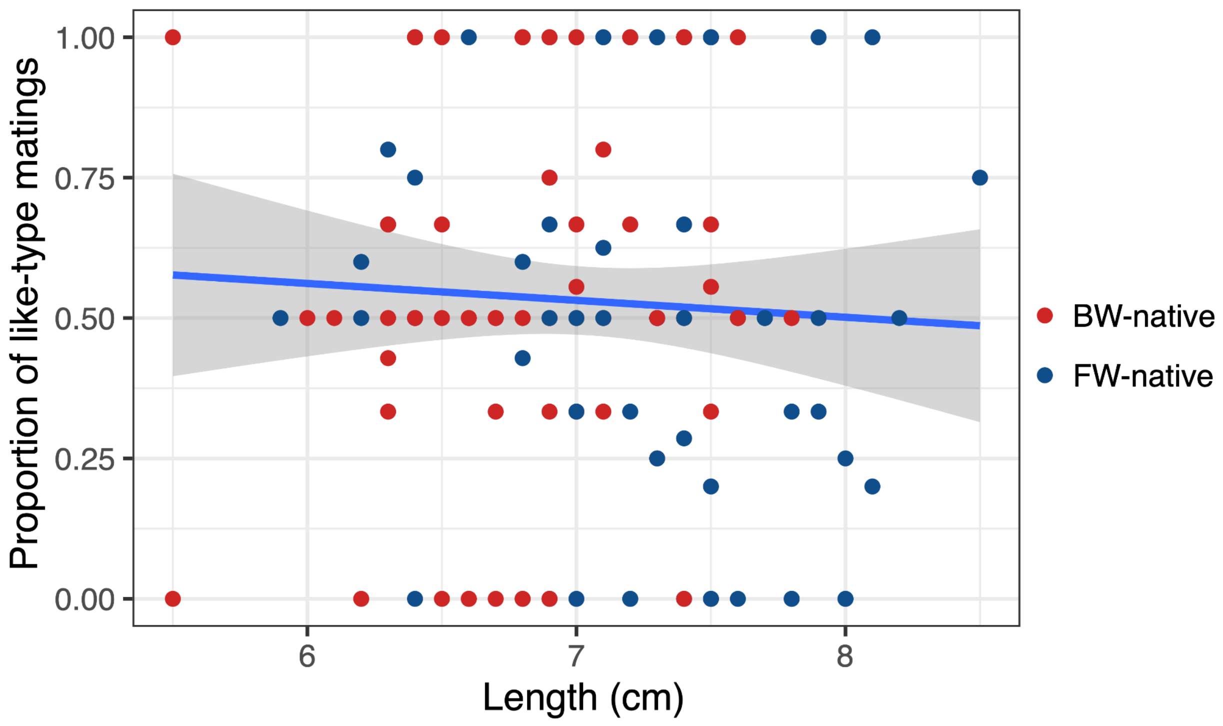


**Figure S8:** Relationship between length and like-type matings. Color indicates the population of each individual.
